# OUGene 2.0: An updated disease-associated over- and under-expressed gene database by mining full-text articles

**DOI:** 10.1101/2022.07.04.498774

**Authors:** Erdi Qin, Xiaoyong Pan, Hong-Bin Shen

## Abstract

Many diseases are closely associated with over- or under-expressed genes. In order to cover more up to date associations between over- or under-expressed genes and various diseases, we develop an updated database OUGENE 2.0 for disease-associated over- and under-expressed genes by automatic full-text mining. In total, the new OUGene 2.0 includes 197,236 associations between 12,672 diseases and 11,542 over- or under-expressed genes, which increases by about 5 folds compared to the previous version of OUGene. A novel method for rescaling the raw score based on support evidences is designed to prioritize the mined associations. OUGene 2.0 provides a holistic view of disease-gene associations and it supports user-friendly data exploration at www.csbio.sjtu.edu.cn/bioinf/OUGene for academic use.

## Introduction

Many diseases are related with various genes’ over- or under-expressions [1]. To date, there exists a huge number of disease-associated over- and under-expressed genes reported in the biomedical literatures. Given a disease, identifying its associated genes by reading through these literatures manually is time-consuming. Thus, it will be useful to automatically extract associations between various diseases and over-and-under-expressed genes from the literatures, to provide a timely access to these data.

The first version of OUGene [2] database extracts the associations between various diseases and over- or under-expressed genes from the abstracts before 2015 in PubMed. Due to the fast growth in the number of PubMed articles, increasing findings on disease-gene associations are reported. Currently there exists some disease-gene association databases, like DISEASES [3] and DisGeNet [4], from text mining and other data sources. However, most of the mined associations are mainly extracted from only the abstracts. In addition, more associations are mentioned in the sentences of the full-text, which provides more disease-gene associations. Automatically extracting the associations from the full-text articles will greatly increase the number and facilitate a global overview of the disease-gene relationships into identifying novel pathways related to diseases [5].

To improve the access to the information on disease-associated over- and under-expressed genes, we update our previous version of OUGene database to OUGene 2.0, with much more number of associations mined from the full-text articles, where the number of mined associations increase from ~ 40,000 to 197,236 with about a 5-fold increase. One primary reason is that OUGene 2.0 mines the associations from full-text article instead of only abstracts. OUGene 2.0 makes full use of text mining to automatically extract associations from full-text articles, and it provides a valuable resource for disease-associated over- and under-expressed genes. The web interface of OUGene 2.0 can be used to analyze these data and visualize the association networks.

We developed a full-text mining-based pipeline to extract associations between diseases and over- and under-expressed genes to construct OUGene 2.0, where RENET2 is used as the text mining engineer. RENET2 [6] is a high-performance full-text gene-disease relation extraction approach, which implements Section Filtering and ambiguous relations modeling with an iterative training data expansion strategy. The whole pipeline of OUGene 2.0 mainly consists of the following four steps (Figure 1): 1) Construction of disease, gene and cell line dictionaries and full-text biomedical articles from PubMed; 2) Using RENET2 to automatically to extract potential associations between over-and-under-expressed genes and diseases; 3) Benchmarking the association against curated data and ranking the mined associations by the designed scoring system; 4) Construction of easy-to-use web interface for querying disease-associated over- and under-expressed genes.

**Figure 1.**
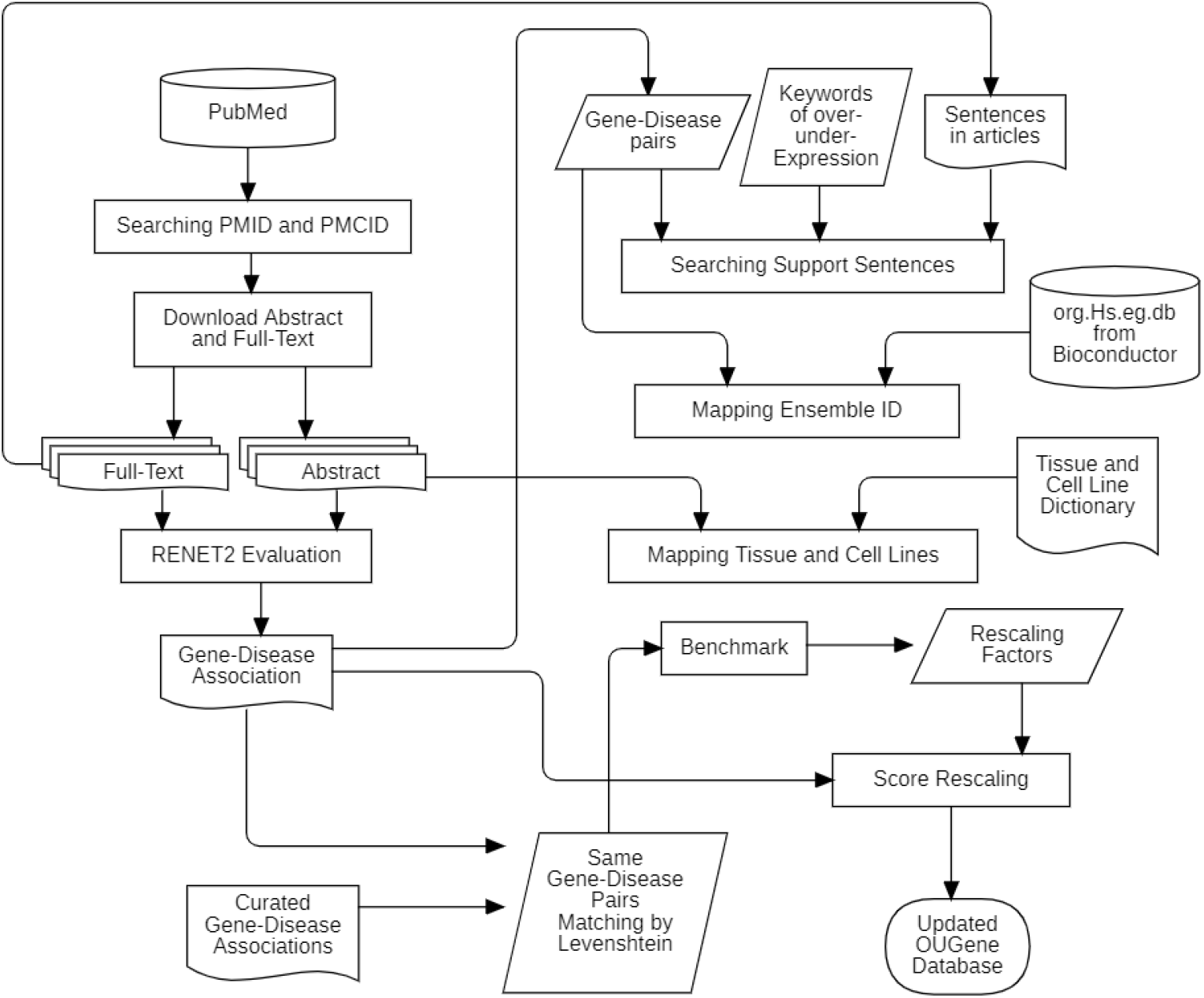
The flowchart of OUGene 2.0 for collecting disease-associated over- and under-expressed genes from full-text articles.

Gene-disease association pairs are obtained from the full-text mining approach RENET2. When analyzing a certain article, if both gene name and disease name appear in the same sentence with the keywords like ‘over’, ‘Over’, ‘under’, ‘Under’, ‘express’, ‘Express’, it is considered as a potential target sentence. During the process, multiple records of gene-disease associations of the same sentence are examined by traversing the full-text once, where the time complexity was o(n) for searching. Then, we obtain the records consisting of PubMed ID, gene name, disease name, expression (over- or under-), score and evidence sentence.

Further, we map gene names to Ensembl name based on the database org.Hs.eg.db from Bioconductor [7]. Next, we extract the tissue and cell line information where the gene is over- or under-expressed. Tissue and cell line dictionary is extracted from TISSUE S database [8] using fuzzy string matching to detect the cell line and tissues mentioned in the article.

To improve the reliability of the association scores, we design a benchmark system and rescaling algorithm, where the benchmark dataset is collected using Curated gene-disease Associations (hereinafter called ‘CURATED’) from DisGeNet database. We extract the same gene-disease pairs from RENET2 and CURATED by examining their similarity, where gene names are required to be exactly the same and disease names are mapped by calculating their Levenshtein Distance, which was set to 0.8 since it needs a sufficient number of samples to eliminate the contingency with high mapping accuracy. By comparing the residual of scores on the same gene-disease pairs from RENET2 and CURATED, we use *β* to describe the imitative effect:

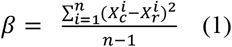

where 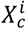 is the association score from CURATED, and 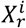 refers to the predicted association score from RENET2 on the same gene-disease pair, *N* is the number of the associations.

When analyzing the distribution of gene-disease association scores from RENET2 and CURATED, we noticed that all the curves approximately fit the normal distribution. In addition, the score difference mainly lies at 0.6 due to the tendency to score higher in RENET2’s scoring system. Meanwhile *β* is approximately 0.20 in the unprocessed data, indicating that score difference lies mainly at 0.5, which may produce misleading information of gene-disease associations. Thus, we develop a rescaling function to adjust the scores of RENET2:

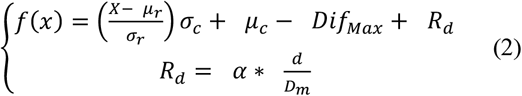

where *X* is the original score from RENET2, *μ_r_*, *σ_r_* are the mean and standard deviation of original scores, *μ_c_*, *σ_c_* is the mean and standard deviation of curated scores, *Dif_Max_* is the translation factor, *R_d_* is the dilation coefficient, *α* is the compensation factor, *D_m_* is the maximum distance on either side of distribution curve peak, *d* is the distance between the input score and the distribution curve peak.

*μ_r_, σ_r_, μ_c_, σ_c_* are used to approximate OUGENE curve to CURATED curve’s patterns, *Dif_Max_* is used to decrease most of the difference under 0.05 by translating the curve directly, since we observe that there is still a deviation after rescaling. *R_d_* is used to reduce the negative effect including the minimum score below 0 brought by the translation, *α* is used to ensure the minimum score is greater than 0, *D_m_* and *d* are used to determine the dilation level depending on the location of input score and the distance from the curve peak, respectively.

For the maximum distance *D_m_*, there are two cases:

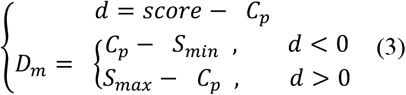

where *C_p_* is the distribution curve peak, *S_min_*, *S_max_* is the minimum and maximum score.

As shown in Figure 2A and B, we can see that the peak difference between original scores and curated scores is about 0.6. After rescaling, we can see that the score distribution between original and curated scores are similar. The results demonstrate that recalling adjust the association scores to be close to the courted scores.

**Figure 2.**
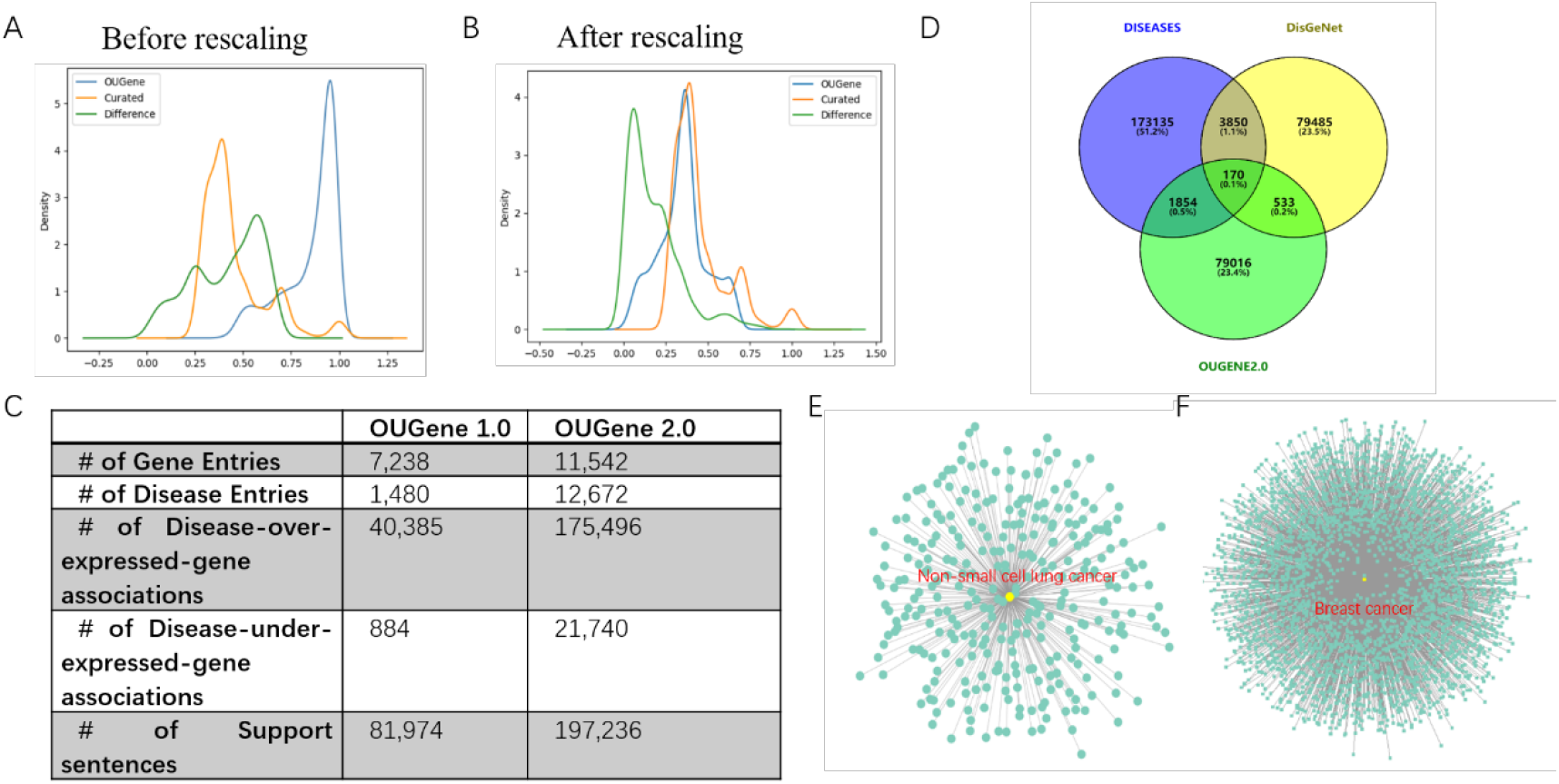
The score rescaling and the statistics in OUGene 1.0 and 2.0. A-B. The score distribution before and after rescaling. C. The statistics in OUGene 1.0 and 2.0. D. Venn diagram of gene-disease associations in DISEASES, DisGeNet and OUGENE2.0. E. The association network between non-small cell lung cancer and over- or under-expressed genes. F. The association network between breast cancer and over- or under-expressed genes.

After rescaling, we calculate the precision, recall and F1-score metrics to evaluate the model performance. Here, we develop an annotating function to define the predicted associations:

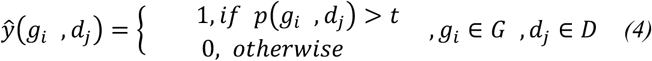

where *t* stands for a threshold, *g_i_*, *d_j_* are gene and disease respectively, *p*(*g_i_*, *d_j_*) means the score predicted by OUGene2.0, G and D is the set of genes and diseases, respectively.

By such an annotating function, we can define true positives, false positives and so on when compared to the CURATED associations from DisGeNet. OUGene 2.0 yields a precision of 0.778, a recall of 0.691 and a F1-score of 0.732 when benchmarked against the curated associations.

As shown in Figure 2C, OUGene 2.0 contains 197,236 associations between 11,542 genes and 12,672 diseases. Of which, 175,496 are the association between disease and over-expressed genes, the remaining 21,740 are associations between diseases and under-expressed genes. We can conclude that most of diseases are associated with gene overexpression [9]. Compared to OUGene 1.0, the number of associations increases from 41,269 to 197,236 (a relative increase about 5 fold), the number of diseases increases from 1,480 to 12,672, and the number of genes increases from 7,238 to 11,542. In addition, the number of support sentences increase about 1.5 fold from 81,974 to 197,236.

As shown in Figure 2D, OUGene 2.0 exploits only a small fraction of gene-disease associations in DISEASES and DisGeNet database, since OUGene 2.0 is focused on gene overexpression or under-expression, but DISEASES and DisGeNet covers all types of associations. We can see that here databases contain different associations between diseases and genes. Furthermore, we illustrate the network between non-small cell lung cancer and its over- and under-expressed genes (Figure 2E). In total, 128 genes are associated with non-small cell lung cancer, of them, eight genes are under-expressed and the remaining genes are over-expressed in non-small cell lung cancer. For breast cancer, it has 1365 associated over- and under-expressed genes (Figure 2F), and the number is much more than that of non-small cell lung cancer

Together with technical improvements on the full-text mining and the increasing published articles, OUGene 2.0’s records have been increased about 5 folds in the number of disease–over-under-under-expressed-gene associations. The database OUGene 2.0 provides an easy-to-use web interface for browsing and searching with network visualization between over-and-under-expressed genes and diseases, which is expected to provide a comprehensive resource for understanding the relationship between gene expression levels and diseases.

